# Nanopore adaptive sampling for mitogenome sequencing and bloodmeal identification in hematophagous insects

**DOI:** 10.1101/2021.11.11.468279

**Authors:** Evan J. Kipp, Laramie L. Lindsey, Cristina M. Blanco, Julia Baker, Marissa S. Milstein, Christopher Faulk, Jonathan D. Oliver, Peter A. Larsen

## Abstract

Blood-feeding insects are important vectors for an array of zoonotic pathogens. Despite significant research focused on well-documented insect vectors of One Health importance, resources for molecular species identification of a large number of hematophagous arthropods are limited. Advancements in next-generation sequencing technologies provide opportunities for targeting mitochondrial genomes of blood-feeding insects, as well as their bloodmeal hosts. This dual approach holds great promise for elucidating complex disease transmission pathways and enhancing the molecular resources for the identification of cryptic insect species. To this end, we leveraged the newly developed Oxford Nanopore Adaptive Sampling (NAS) pipeline to dually sequence the mitogenomes of hematophagous insects and their bloodmeals. Using NAS, we sequenced the entire mitogenomes of *Aedes vexans*, *Culex restuans*, *Culex territans*, and *Chrysops niger* and successfully identified bloodmeal hosts of *Chrysops niger*, *Culex restuans*, and *Aedes trivittatus*. We show that NAS has the utility to simultaneously molecularly identify blood-feeding insects and characterize disease transmission pathways through bloodmeal host identification. Moreover, our data indicate NAS can facilitate a wide array of molecular systematic studies through novel ‘phylogenetic capture’ methods. We conclude the NAS approach has great potential for informing global One Health initiatives centered on the mitigation of vector-borne disease through dual vector and bloodmeal identification.

## 1. Introduction

Hematophagous insects are major disease vectors that transmit a wide variety of pathogens to their bloodmeal hosts. From a One Health perspective, blood-feeding insects are responsible for considerable morbidity and mortality across global human, livestock, and wildlife communities (Pérez de León, Mitchell, & Watson, 2020). Yet, despite their global importance, these diseases remain difficult to study and adequately control. Vector-borne pathogens are often maintained in complex enzootic transmission cycles which frequently involve numerous species of both vector and vertebrate hosts. Even among groups of hematophagous insects (e.g., mosquitoes) there exists a wide diversity of distinct species which may individually exhibit substantial differences in their host feeding preferences, vectorial capacity, and other natural history characteristics which impact their ability to serve as disease vectors (Gómez-Díaz & Figuerola, 2010; Kent, 2009). Fortunately, the advancement of molecular and genetic tools throughout the past decades has helped to inform many complex aspects of vector-borne pathogen transmission. DNA barcoding—accomplished through PCR amplification and sequencing of select phylogenetically informative loci—has facilitated species identification efforts across hematophagous insect taxa (Beebe, 2018; Jinbo, Kato, & Ito, 2011), and has been successfully applied to the molecular identification of vertebrate hosts from arthropod bloodmeals (Alcaide et al., 2009; Mukabana, Takken, & Knols, 2002; Reeves, Gillett-Kaufman, Kawahara, & Kaufman, 2018; Townzen, Brower, & Judd, 2008).

Across metazoan organisms, mitochondrial genes have emerged as the most popular targets for such species barcoding efforts. In comparison to most nuclear genes, those encoded on mitochondrial genomes (mitogenomes) are well-suited for molecular species identification and phylogenetic reconstruction due to their conserved evolutionary origins, indispensable cellular functions, uniparental inheritance, and elevated mutation rates combined with very low rates of recombination (Gissi, Iannelli, & Pesole, 2008). Animal mitogenomes are traditionally circular in structure, relatively small in size (~16kb), and contain the same core set of 13 protein-coding genes (PCGs), 22 transfer RNAs (tRNAs), and 2 ribosomal RNAs (rRNAs)(Boore, 1999; Gissi et al., 2008). Furthermore, mitogenomes are present at many copies within each cell, making them convenient targets for molecular analyses using a variety of biological sample types (Boore, 1999; Cameron, 2014; Crampton-Platt et al., 2015; Crampton-Platt, Yu, Zhou, & Vogler, 2016; Romero, Weigand, & Pfenninger, 2016; Timmermans et al., 2010; Wanner, Larsen, McLain, & Faulk, 2021). Sequence data of numerous mitochondrial loci (e.g., *cytochrome c oxidase subunit 1* (COI or cox1), *cytochrome b* (cyt-b), and D-Loop) are well-documented for their phylogenetic and phylogeographic utility, having variable rates of evolution that can be leveraged to test taxonomic hypotheses and elucidate evolutionary histories (Beebe, 2018; Bradley & Baker, 2001; Creedy et al., 2021; Jinbo et al., 2011; Townzen et al., 2008). To date, the majority of barcoding efforts, including those focused on blood-feeding insects and bloodmeal analysis, have leveraged traditional PCR followed by Sanger sequencing of either COI and cyt-b (Beebe, 2018; Borland & Kading, 2021; Jinbo et al., 2011; Molaei et al., 2007; Santiago-Alarcon, Havelka, Schaefer, & Segelbacher, 2012; Schnell et al., 2012; Townzen et al., 2008; Videvall et al., 2013). Although such molecular barcoding has proved useful for identifying particular taxonomic lineages, it remains limited in its generation of singular-gene sequence data and potential for false-negative results due to PCR failure.

The weaknesses of PCR-based species barcoding and bloodmeal analysis may be largely overcome using *de novo* next-generation metagenomic sequencing approaches. Previously, high-throughput applications (i.e., Illumina sequencing) have been shown to provide ample depth-of-coverage required for the *de novo* taxonomic classification of bloodmeal hosts (Borland & Kading, 2021; Kieran et al., 2017; Muturi, Dunlap, Tchouassi, & Swanson, 2021). Despite technological and bioinformatic advancements, second-generation sequencing methods are time-consuming, frequently require large brick-and-mortar sequencing laboratories, and cannot be performed in diverse field settings. When considering the growing burden of vector-borne diseases in developing nations, especially across tropical regions of the world, second-generation metagenomic approaches are largely out of reach due to limited availability.

Third-generation nanopore sequencing technologies and associated bioinformatics have opened the door to a variety of global One Health research initiatives. The Oxford Nanopore Technologies (ONT) MinION sequencing platform is a portable device and thus can be deployed globally, including in remote field locations (Blanco et al., 2020; Hoenen et al., 2016; Jain, Olsen, Paten, & Akeson, 2016). The MinION requires basic molecular lab equipment and downstream bioinformatic approaches can be performed locally on either a desktop or laptop computer. An important aspect of the MinION platform is that it sequences individual DNA or RNA molecules across a microfluidic flowcell. During a MinION experiment, single-molecule nucleotide sequences of the sequencing library are typically produced at a rate of ~450 bases per second; hundreds of DNA/RNA fragments are analyzed at once. Given the novelty of nanopore-based real-time sequencing, a variety of bioinformatic pipelines have been developed that effectively leverage the single-molecule aspect of the technology.

Nanopore adaptive sampling (NAS) is a recently developed method that selectively sequences individual DNA, cDNA, or RNA molecules in real-time using ONT sequencing technology (Kipp et al., 2021; Payne et al., 2021; Wanner et al., 2021). NAS uses an advanced bioinformatic pipeline that compares nucleotides of individual molecules (~200-400 base pairs) against a reference file every ~0.4 sec, as sequencing is occurring. This dynamic method has a wide number of applications and can selectively enrich DNA or RNA targets of interest (e.g., particular genes, RNA species, mitogenomes, etc.) and reject unwanted molecules during a sequencing experiment (Kipp et al., 2021; Payne et al., 2021). Here, we show how the NAS method can be leveraged to target and sequence entire mitochondrial DNA genomes (mitogenomes) of hematophagous insect vectors. Additionally, we show proof-of-concept for bloodmeal host identification using the NAS method (Fig 1). NAS, combined with the long-read sequencing capabilities of the ONT MinION, is ideally suited for the selective enrichment and assembly of complete mitogenomes (Wanner et al., 2021). Moreover, NAS mitogenome sequencing can effectively capture mitochondrial sequences having at least 75% identity with a given reference; thus, the method can be used for mitogenome assembly of a variety of taxa for which references are absent (Wanner et al., 2021). We used the NAS mitogenome sequencing approach to molecularly characterize four mosquito species as well as a biting deer fly and its bloodmeal host from Minnesota. We posit that NAS holds great promise for a variety of One Health research projects aimed at elucidating hematophagous insect diversity and associated pathogen transmission pathways.

**Figure 1.**
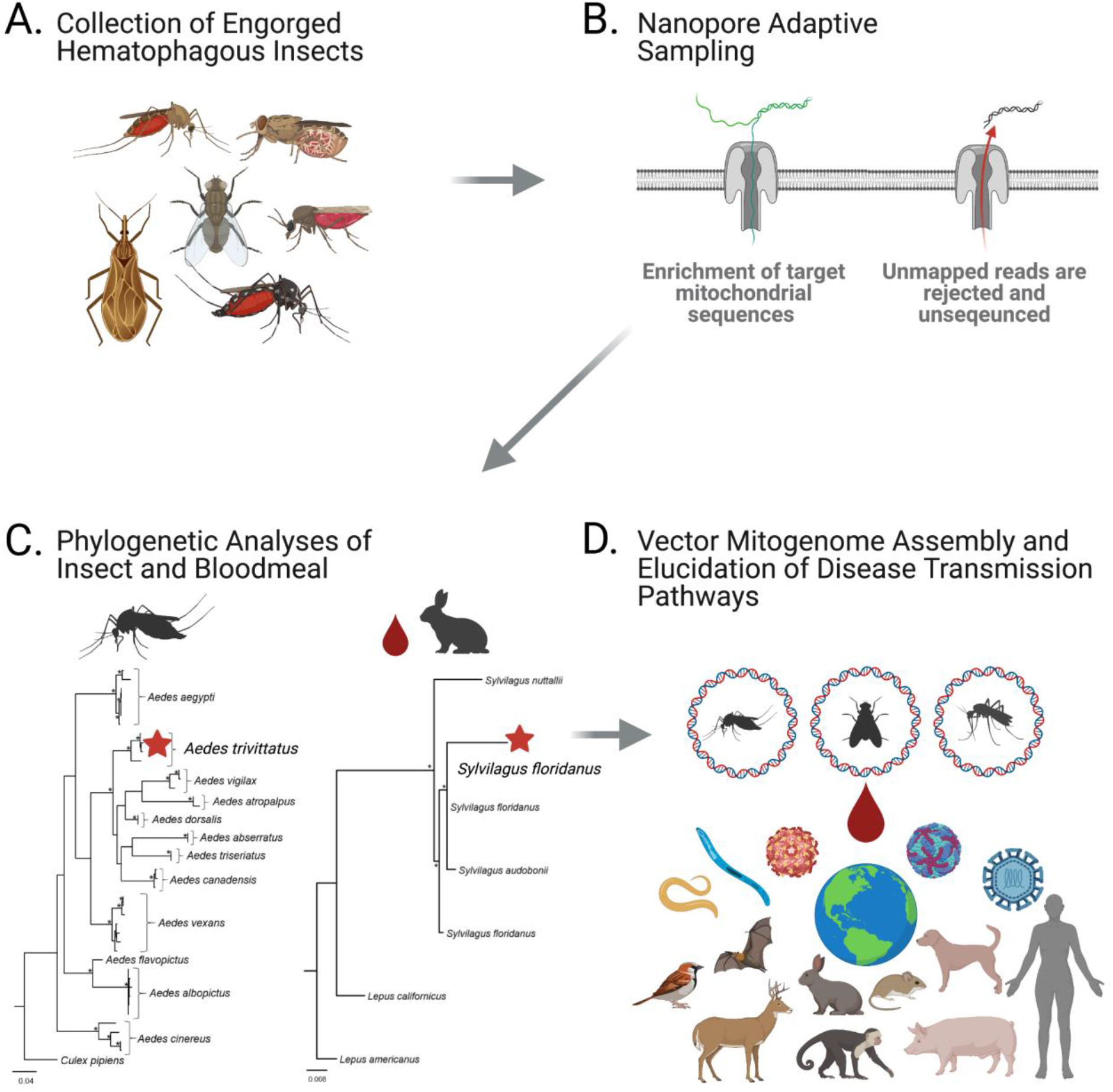
Experimental design for dual insect and bloodmeal mitochondrial DNA sequencing using nanopore adaptive sampling. **A:** Hematophagous insects having recent bloodmeals are collected from natural populations. Whole genomic DNA is extracted from individual or pooled insects. **B:** Genomic DNA is prepared for sequencing on an Oxford Nanopore Technologies device with access to the Nanopore Adaptive Sequencing (NAS) bioinformatic pipeline. NAS is performed on blood-feeding insect DNAs with a reference file containing mitochondrial DNA sequences of congeneric or conspecific insect species and a diverse selection of putative bloodmeal hosts. Exact species matches are not needed for NAS references as the method will retain sequences sharing at least 75% identity, thus suitable for guided discovery of cryptic insects and bloodmeal hosts. **C:** Mitochondrial sequences from NAS vector and host matches are quality filtered and species barcoding genes (i.e., COI, cyt-b, D-Loop, etc.) are used for phylogenetic analyses and species identification. **D:** Complete mitogenomes are recovered from the sampled insect, thus expanding molecular resources for vector species. Recovered mitochondrial sequences of bloodmeal hosts elucidate potential disease transmission pathways. Collective results of NAS experiments directly inform One Health initiatives focused on hematophagous insect biology and vector-borne disease transmission.

## 2. Material and methods

### 2.1 Specimens examined

Mosquitoes were collected from a forested area and a residential area in Ramsey County, Minnesota within the Minneapolis-St. Paul metropolitan region using dry ice-baited CDC Miniature Light Traps (model 512; John W. Hock Company; Gainesville, United States) hung approximately six feet above ground. Traps were deployed overnight from approximately 17:00hrs to 10:00hrs the following day. A single bloodfed *Chrysops* deer fly was opportunistically collected while conducting fieldwork within Washington County, Minnesota (Afton State Park; as approved by the Minnesota Department of Natural Resources: permit 202145). All specimens were cold anesthetized and morphologically examined under a stereomicroscope. Bloodfed female mosquitoes were separated from non-target insects and initial taxonomic identifications were based on standard morphological features (Darsie & Ward, 2005). All specimens were submerged in RNAlater (Sigma Aldrich, St. Louis, United States) and subsequently preserved at −80 °C prior to nucleic acid extraction and molecular analyses. A list of specimens examined is provided in Supplemental Table 1.

### 2.2 DNA extraction and ONT library preparation

Genomic DNA was individually extracted from bloodfed mosquitoes of three species: (*Cu. restuans (*n= 3), *Cu. territans* (n= 1), *Ae. vexans* (n= 1), and *Ae. trivittatus* (n=1)); and one species of deer fly (*Ch. niger* (n= 1)), using a Qiagen DNeasy Blood and Tissue Kit following manufacturer’s instructions (Qiagen, Hilden, Germany). Resulting extracts were quantified using a Qubit 4 Fluorometer (Invitrogen, Waltham, United States). Genomic DNA libraries for ONT sequencing were produced for each specimen using Sequencing Ligation Kits SQK-LSK109 and SQK-LSK110, following standard ONT protocol. For each sample, 0.3 to 1.5 μg of initial DNA was used for ONT sequencing library construction following Kipp et al. (In review). Samples were barcoded using ONT Kit EXP-NBD104, pooled, and five libraries were sequenced until completion for ~22-48 hours on R9.4 flowcells. All sequencing was performed on a Linux desktop computer with the following specifications: Intel C600/X79 series i9-10920X 12 core; Linux 5.4.0-77-generic x86_64; Ubuntu 18.04; Nvidia Quadro RTX 4000 GPU. The 5 sequencing experiments are denoted herein as follows: A: *Ch. niger* NAS (2 barcodes - 1 barcoded sample consisting of midges not included in the present study); B: *Cu. restuans* and *Cu. territans* barcoded NAS; C: *Ae. vexans* NAS (no barcode); D: *Cu. restuans* unenriched control sequencing (no barcode and no NAS); E: *Cu. restuans* and *Ae. trivittatus* barcoded NAS (2 additional barcoded samples not included in the present study) (see Table 1)

**Table 1.** Nanopore sequencing runs conducted to sequence genomic DNA from 5 vector species containing bloodmeals. User-generated reference files are included in Supplemental Table 2. The number of bases and reads generated for each sample and average read lengths (N50) are reported below. (*Specimens from these runs were used for whole insect mitogenome assemblies).

| Experiment ID | NAS technique | Sample | Run time | Total bases | # Reads (million) | Avg read length (N50) in bp per experiment |
| --- | --- | --- | --- | --- | --- | --- |
| A* | Enriched using user-generated reference file | *Chrysops niger* | 22h 42m | 600 Mb | 1.5 | 401 |
| B* | Enriched using user-generated reference file | *Culex restuans* | 45h 47m | 1 Gb | 2.5 | 462 |
| B* | Enriched using user-generated reference file | *Culex territans* | 45h 47m | 2 Gb | 4.5 | 462 |
| C* | Enriched using mitogenome database as reference | *Aedes vexans* | 21h 7m | 4 Gb | 10.8 | 536 |
| D | No NAS. Control run. | *Culex restuans* | 44h 58m | 5 Gb | 6.2 | 1,960 |
| E | Enriched using user-generated reference file | *Culex restuans* | 27h 15m | 350 Mb | 1.0 | 347 |
| E | Enriched using user-generated reference file | *Aedes trivittatus* | 27h 15m | 200 Mb | 0.6 | 347 |

### 2.3 Basecalling

Following Kipp et al. (In Revision), sequencing was initiated using the MinKNOW GUI, v4.3.20, in which adaptive sampling software is integrated. Reference files containing publicly available mitogenome sequences and sequences of select barcoding genes (i.e., COI, cyt-b) were compiled and provided to the MinKNOW for NAS-based enrichment (GenBank accession numbers are provided in Supplemental Table 2). Real-time basecalling was performed using the fast basecalling model in Guppy and mapped to a provided reference file (see Supplemental Table 2 for a detailed list of each reference file) using the NAS MinKNOW software. After completion of sequencing runs, post-hoc high accuracy basecalling was performed using Guppy (v5.0.11; release with GPU-enabled basecalling for Ubuntu) prior to downstream analyses. The NAS software pipeline provides an optional immediate readout of reads mapping to particular reference sequences, and real-time alignment data can be recorded in the resulting sequencing summary output text file. For Experiments C and E, this optional readout was utilized to extract individual sequences based on successfully aligned reads contained in the sequencing summary output text file. For the remaining NAS experiments (A and B), targeted mitochondrial reads were mapped using minimap2 (v2.17-r941) and indexed using samtools (v1.9) in post-hoc bioinformatic analyses.

### 2.4 NAS versus control sequencing

A control nanopore sequencing experiment (D) was conducted to test the effectiveness of NAS. This experiment generated sequences from genomic DNA of a single mosquito (*Cu. restuans*) without NAS for nearly 45 hours. During the sequencing run, the basecalling was set to fast mode (similar to NAS protocols) and post-hoc basecalling was conducted using the high accuracy mode of Guppy to keep all other sequencing variables (other than NAS) the same. Sequences from Experiment D were filtered based on a Qscore of 10 and read lengths between 1kb and 16kb using NanoFilt v2.7.1 (De Coster, D’Hert, Schultz, Cruts, & Van Broeckhoven, 2018). Filtered reads were subsampled at random using seqtk v1.3 (https://github.com/lh3/seqtk) and used in comparison analyses. Randomly sampled reads were mapped to the *Cu. pipiens* mitochondrial genome using the program minimap2 v2.17 (Li, 2018) and the percent of assembled reads were noted. Experiment D was compared to NAS Experiment B for *Cu. restuans* and *Cu. territans*, using the same filter parameters, random subsampling technique, and mapping program.

### 2.5 Mitogenome assemblies and phylogenetics

Reads were filtered for mitogenome assembly using NanoFilt set to a minimum Q-score of 10 and read lengths between 1kb and 16kb (De Coster et al., 2018). Filtered reads for individual samples were *de novo* assembled using Flye v2.8.3 software following Wanner et al. (2021). Subsequent mitogenome assemblies were polished to produce more accurate assemblies using Medaka v1.4.3 (https://github.com/nanoporetech/medaka). Open reading frames (ORFs) were identified and annotated using MITOS2 web server (Donath et al., 2019; http://mitos2.bioinf.uni-leipzig.de/index.py) with specification for metazoan RefSeq and invertebrate genetic code. Each annotated mitogenome was visually inspected using Geneious Prime software v2021.2.2, and aligned against a published and annotated mitogenome of another member in each genus. Where necessary, the start and end positions of particular genes were manually adjusted based on those of previously-characterized mitogenomes. Polished and assembled mitogenomes were visualized using OGDRAW v1.3.1 (Greiner, Lehwark, & Bock, 2019; https://chlorobox.mpimp-golm.mpg.de/OGDraw.html).

After annotation of the mitochondrial genomes, COI sequences were used in phylogenetic analyses to verify species identification. COI sequences were downloaded from GenBank for each appropriate species (*Culex*, *Aedes*, and *Chrysops*) to build phylogenies. Appropriate outgroup samples also were downloaded from GenBank and included in each analysis to root the phylogeny. Sequences were aligned using MAFFT v7.475 (Katoh & Standley, 2013), and subsequent alignments were used in a maximum likelihood analysis with the following parameters in RAxML v8.211 (Stamatakis, 2014): 1) model of evolution was set to GTRGAMMAI and 2) analyses were performed using 1,000 bootstraps. The resulting best tree for each analysis was visualized using FigTree v1.4.1 (https://github.com/rambaut/figtree).

Bloodmeal identification was determined by mapping reads to the mitochondrial genomes (*Homo sapiens* and *Passer domesticus*) or mitochondrial 12S gene (*Sylvilagus floridanus*) of the respective species. Average coverage was calculated for the human bloodmeal; and phylogenetic trees were generated (using the same parameters mentioned above) for the house sparrow and eastern cottontail bloodmeals to determine statistical nodal support because coverage was significantly lower for these two bloodmeals.

## 3. Results

### 3.1 Output and performance of nanopore sequencing experiments

Genomic DNA was successfully isolated from seven bloodfed insect specimens: *Ch. niger* (1), *Cu. restuans* (3), *Cu. territans* (1), *Ae. vexans* (1), and *Ae. trivittatus* (1). Molecular data for each specimen was generated across five ONT sequencing runs (Table 1). The number of bases generated for each sample ranged from 200 Mb to 5 Gb, and the number of reads generated ranged from 0.6 million to 10.8 million reads. Average read length (N50) was lower in sequences generated from NAS runs (347 - 536bp) compared to the control experiment (1.96 kb), as expected (Table 1). Raw fastq files generated for each sequencing run were submitted to the Sequence Read Archive (SRA) at NCBI (accession numbers reported in Supplemental Table 1).

### 3.2 Comparison of NAS and control sequencing for mitogenome enrichment

Our comparison of the NAS method (Experiment B) versus a control nanopore sequencing experiment (i.e., NAS pipeline off; Exp. D) revealed that approximately 65.8% and 75% of reads in the NAS active experiment (Exp. B: *Cu*. *territans* and *Cu*. *restuans*, respectively) mapped to the *Cu. pipiens* reference, whereas 1.8% of reads in the control experiment (Exp. D) mapped to the same reference (Table 2). Given Exp. D consisted of a single mosquito, we performed sequence subsampling analyses and found similar percentages (NAS: >65% reads mapped; control: ~2% reads mapped; Supplemental Table 3).

**Table 2.** Percentage of total reads mapping to a *Cu. pipiens* mitochondrial genome reference for NAS versus control sequencing experiments. Analyzed samples consisted of a single *Cu. restuans* within the control ONT sequencing experiment (Exp. D) and 2 barcoded samples (*Cu. restuans* and *Cu. territans*) from an NAS-active experiment (Exp. B). Samples were filtered based on a Qscore of 10 and read lengths ranging between 1kb-16kb. The mitogenome of *Cu. pipiens* (NC_015079) was used as a reference for mapping using minimap2 software.

| Experiment ID | Sample | # Seqs after filtering | # Seqs mapped | % Reads mapped |
| --- | --- | --- | --- | --- |
| D | *Culex restuans* | 1,609,492 | 29,289 | 1.8 |
| B | *Culex restuans* | 9,664 | 7,247 | 75.0 |
| B | *Culex territans* | 18,736 | 12,299 | 65.6 |

### 3.3 Assembly and annotation of mitochondrial genomes

For each species sequenced using the NAS enrichment approach, we assembled a single, circular contig from aligned mitochondrial reads. Mean assembly coverage for each mitogenome is reported as follows: *Ch. niger:* 56x, *Cu. restuans*: 612x, *Cu. territans*: 1,342x and *Ae. vexans*: 750x. In addition to these successfully assembled mitogenomes, we also attempted assembly using aligned reads generated during the control experiment (Exp. D: *Cu. restuans*). Here, we noted that 19 contigs of high coverage (1,846x - 2,428x) and short lengths (426bp - 958bp) were generated during Flye assembly and this ultimately precluded our ability to obtain a high-quality mitogenome in the absence of NAS-based enrichment. The four NAS-derived mitogenomes were all of an intermediate size, ranging from 15,573 to 16,048 bp in total length (Figure 2). Nucleotide bias was also consistent across all four mitogenomes and ranged from 78.1% A+T to 79.0% A+T in total nucleotide content.

**Figure 2.**
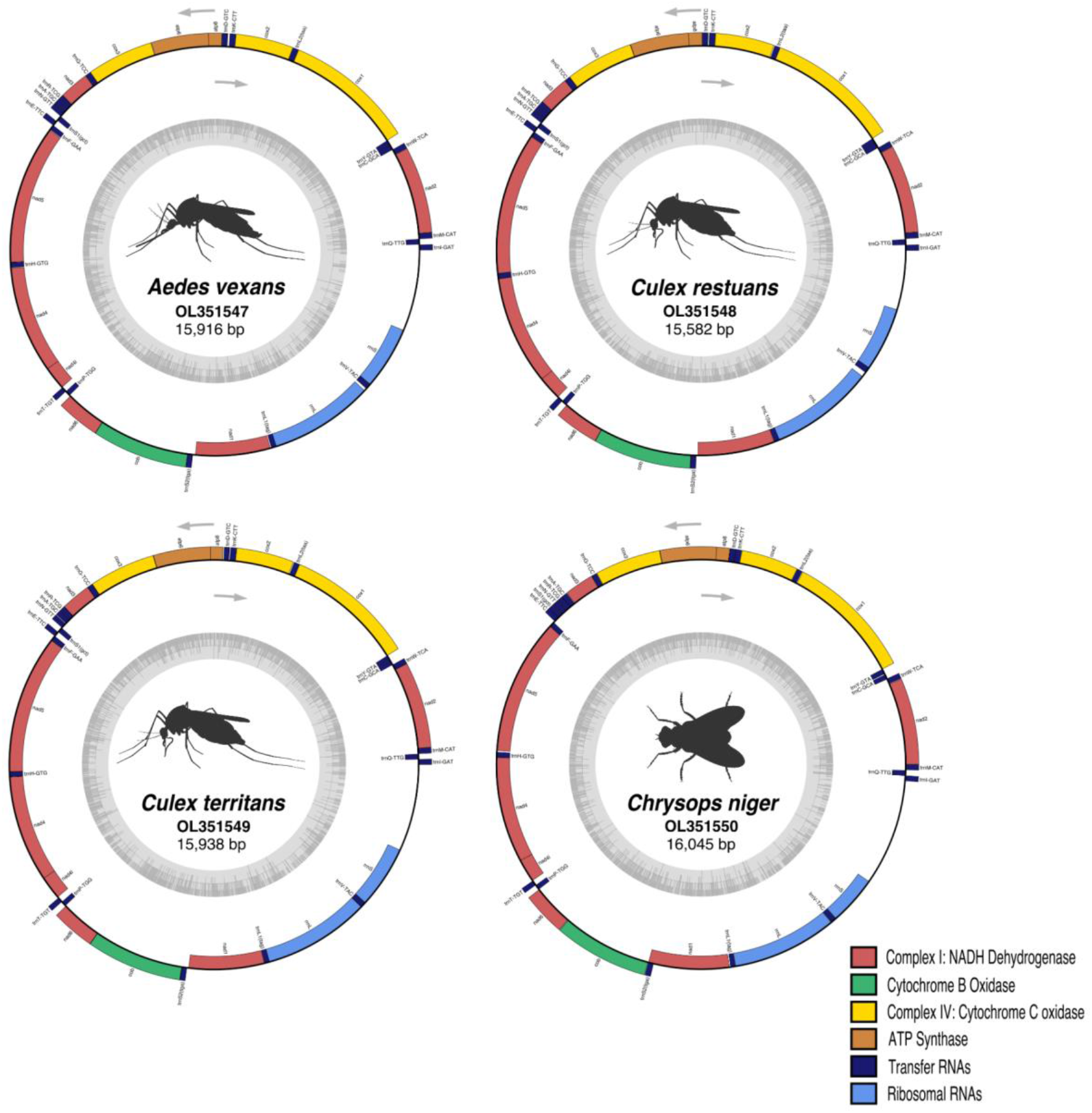
Circular mitochondrial genome maps of the four hematophagous insects sequenced using NAS. Orientation of gene transcription is indicated with gray arrows for genes encoded on the majority, J-strand (counterclockwise, outer circle) and minority, N-strand (clockwise, inner circle). Relative A+T content is visualized as a percentage on the innermost circle in light gray.

Annotation of our assemblies indicated that each encoded a total of 37 genes, which included the expected assemblage of PCGs (13), tRNAs (22), and rRNAs (2). A short region lacking gene content (i.e., control region) at the suspected site of replication initiation was also observed in each sequenced mitogenome. Among our *Aedes* and *Culex* mitogenomes, the majority strand (J-strand) was found to encode nine PCGs and 13 tRNAs, while the minority strand (N-strand) was found to encode four PCGs, nine tRNAs, and both rRNA genes. Overall, this organization of gene content as well as the specific order of genes was found to be identical among our three mosquito mitogenomes and consistent with that reported in other members of the Culicidae family (Behura et al., 2011; Luo et al., 2016). Our *Ch. niger* deer fly mitogenome differed slightly in that the J-strand encoded 9 PCGs and 14 tRNAs, with the N-strand encoding the remaining four PCGs, eight tRNAs, and both rRNAs. Importantly, this organization is consistent with previously-sequenced mitogenome for dipteran members of the suborder Brachycera (Wang et al., 2016). Complete annotated mitogenome assemblies were submitted to the NCBI Organelle database (accession numbers are reported in Supplemental Table 1).

### 3.4 Molecular identification of insect species and bloodmeals

Genetic identification for each vector species was determined using the complete COI gene that was annotated from the assembled mitogenomes reported herein (Experiments A-C) or from consensus sequences generated by mapping to a COI reference (Experiments D and E). The resulting best trees generated from a bootstrapped maximum likelihood analysis are reported in Supplemental Figure 1. Each vector COI sequence generated for this study was statistically supported and formed a monophyletic clade of the appropriate species.

A bloodmeal identified as human was successfully sequenced from *Ch. niger* (Exp. A). We recovered over 200 mitochondrial sequences of the human bloodmeal and mapped reads primarily ranged from ~100 bp to ~8kkp in length, with a single read spanning nearly the entire human mitochondrial genome (16,123bp; Figure 3). Experiment E generated bloodmeal sequencing reads for two vector species. For the *Cu. restuans* vector, 16 reads mapped to the house sparrow (*Passer domesticus*) mitochondrial genome. Mapped reads had a range of 106 base pairs to 2,475 base pairs in length. Of these 16 mapped reads, 3 reads mapped to the control region (CR) of the house sparrow mitochondrial genome. A maximum likelihood analysis revealed the sequences generated from the CR formed a statistically supported clade with *Passer domesticus* (1.0% divergence within the *P*. *domesticus* clade; Fig 4A). The *Ae. trivittatus* vector from Experiment E resulted in a bloodmeal host identified as an eastern cottontail rabbit (*Sylvilagus floridanus*) with a single read (955 bases long) mapping to the 12S gene. A phylogenetic analysis of the 12S gene supports the molecular identification of the bloodmeal to *Sylvilagus floridanus* (2.0% divergence within the *S*. *floridanus* clade; Fig 4B).

**Figure 3.**
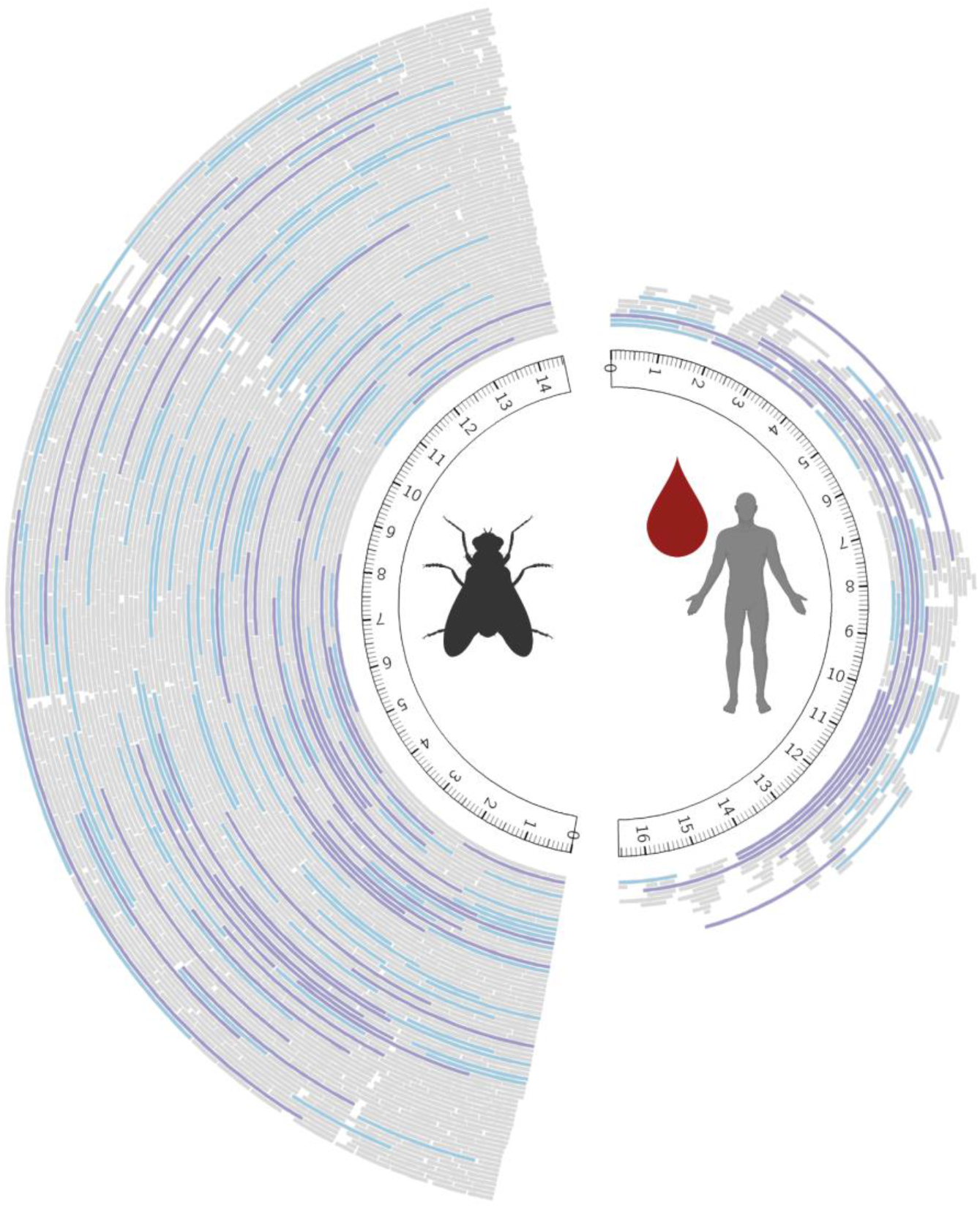
Circos diagram showing NAS reference-guided mitogenome mapping results for a Minnesota black deer fly *Ch. niger* (left; reference *Ch. silvifacies* from China NCBI KT225292.1) and its human bloodmeal (right; reference human mitogenome NCBI NC_012920.1). Outer layers depict individual ONT reads mapping to reference mitogenomes where >2kb reads are identified in purple, 1-2kb in blue, and <1kb are gray. Reference mitogenomes for *Ch. silvifacies* and human are identified on the left and right, respectively, with labeled major tick marks every 1kb and unlabeled minor ticks every 100bp.

**Figure 4.**
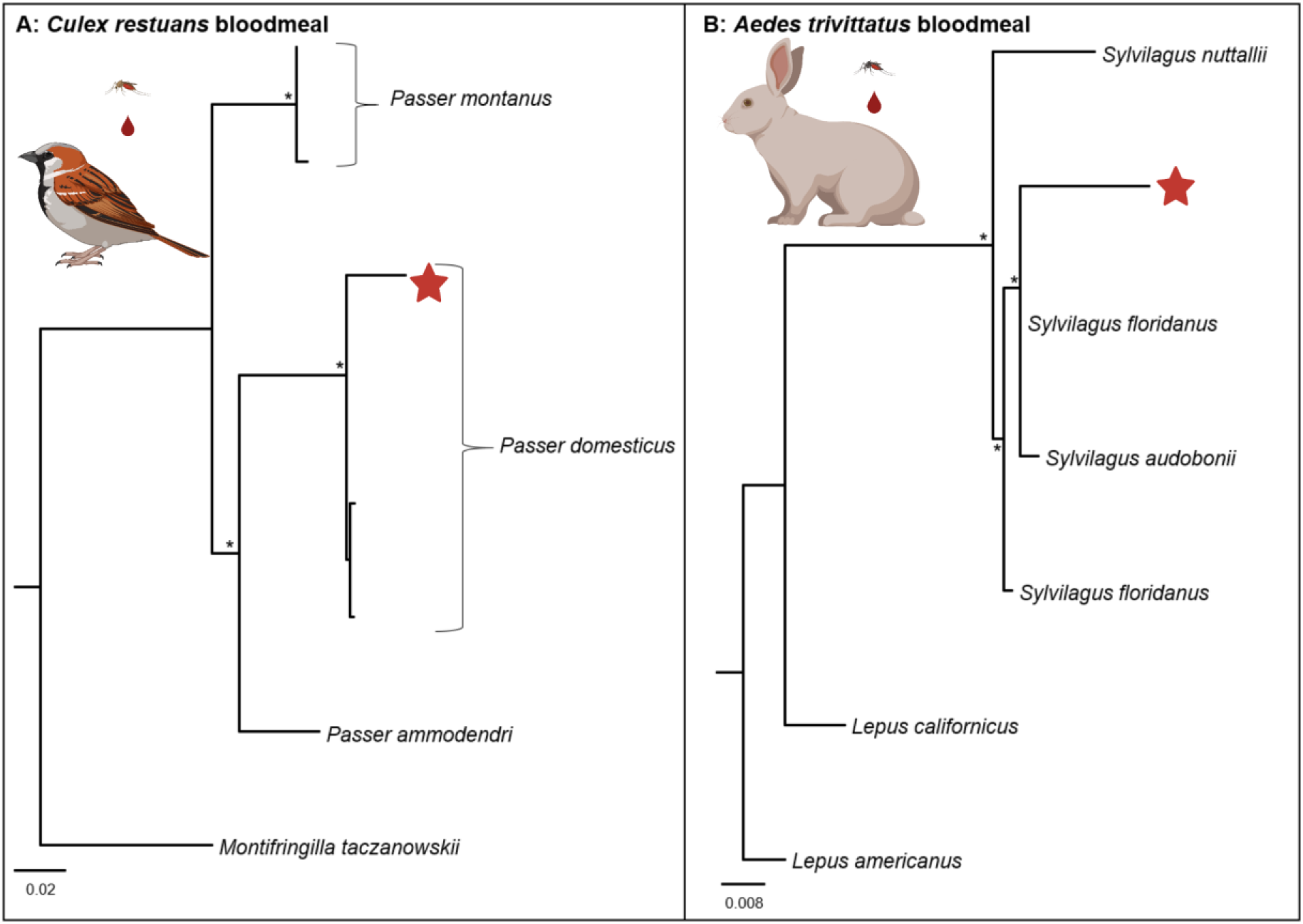
Phylogenetic trees constructed of *Culex restuans* and *Aedes trivittatus* bloodmeals. Barcoded sequences generated from *Cu. restuans* and *Ae. trivittatus* (Exp. E) are denoted by a red star and statistically supported nodes (≥ 75) are denoted with a single asterisk (*). Genbank accession numbers of sequences included in both phylogenetic analyses are provided in Supplemental Table 4. **A:** Maximum likelihood analysis with 1,000 bootstrap replicates for the control region (i.e., D-loop) of Passerine birds. **B:** Maximum likelihood analysis with 1,000 bootstrap replicates for the 12S gene of rabbits (genera *Lepus* and *Sylvilagus*).

## 4. Discussion

### 4.1 Hematophagous insect and bloodmeal mitogenome recovery

Using a purely bioinformatic approach, we successfully enriched, sequenced, and assembled entire mitochondrial genomes of three mosquito species *Ae. vexans*, *Cu. restuans*, *Cu. territans*, and the black deer fly (*Ch. niger*) (Figure 2). Bloodmeal identifications were successful for *Cu. restuans* (house sparrow), *Ae. trivittatus* (eastern cottontail rabbit) and *Ch. niger* (human) (Figs 3 and 4; Experiments A and E). However, we were unable to recover bloodmeal identifications for mosquitoes sampled in NAS Experiments B and C. As reviewed by Kent (2009), there are several variables which either individually, or combined, could influence successful bloodmeal identification using molecular methods including: time between feeding event and sample preservation (longer times result in more host blood being digested); mammalian hosts having enucleated red blood cells (successful ID dependent upon leukocytes present in a mammalian sample; increased chance of success if host has nucleated RBCs [e.g., birds, reptiles, amphibians]); and a high percentage of hematophagous insect DNA within extracts (bulk, whole individual, DNA extracts result in majority of DNA originating from the hematophagous insect). Despite these observations, NAS for bloodmeal identification has several advantages over traditional PCR approaches, in particular NAS is not confounded by PCR inhibitors (i.e., heme) present in blood (Akane, Matsubara, Nakamura, Takahashi, & Kimura, 1994) and the method could detect multiple hosts within a heterogeneous bloodmeal when using a reference file of substantial phylogenetic breadth. Moreover, because nanopore sequencing is a third-generation single-molecule technology, NAS data negate the need for cloning and Sanger sequencing of mixed, multi-host, PCR products that are the result of PCR primers targeting highly conserved regions.

### 4.2 Phylogenetic capture

Regarding our NAS-based recovery of the mitogenomes of the *Ch. niger* deer fly, the only available mitochondrial genomes for reference was from *Ch. silvifacies* collected from China. Our *de novo* assembly of the *Ch. niger* mitogenome is separated by a genetic distance of ~9.4% (Kimura-2 Parameter values; Bradley & Baker, 2001) from *Ch. silvifacies.* Likewise, we used the mitochondrial genome for *Cu. pipiens* collected from Tunisia as reference for our *Cu*. *restuans* and *Cu*. *territans* NAS sequencing. The *de novo* assembled mitogenomes generated herein for *Cu*. *restuans* and *Cu*. *territans* have a genetic distance of 5.5% and 8.0% from *Cu*. *pipiens*, respectively. These results indicate NAS can be leveraged with a ‘phylogenetic capture’ approach using distantly related, yet congeneric species. The broader implications of these findings is that NAS-based recovery of mitochondrial DNA sequences requires only a single mitochondrial genome from a given genus. It is also possible that strategically selecting reference sequences for NAS that effectively span the phylogenetic distance of a given clade (i.e., basal and terminal clades, one or more members of a polytomy, one of two sister taxa, etc.) will enhance the discovery and identification of taxa wherein reference sequence data are absent or unavailable. We note that the NAS reference file for Experiment C (*Ae. vexans*) included all publicly available mitogenomes on the NCBI organelle database at the time of our experiment (11,982 mitogenome sequences; accessed on 16 September 2021). We observed no indication that the MinKNOW software or NAS method was impacted by this number of reference sequences. The implication of this observation is that a single mitogenome reference file spanning the tree of life can be used for the NAS phylogenetic capture method. If accurate, this approach has far-reaching implications for species barcoding and molecular systematics that extend beyond the scope of the analyses presented herein.

### 4.3 NAS performance versus control sequencing

Nanopore adaptive sequencing is a relatively new bioinformatic pipeline (Payne et al. 2021), thus comparisons to traditional nanopore sequencing runs are essential. The analyses presented herein and those of others (Gan et al., 2021; Kipp et al., 2021; Martin et al., 2021; Payne et al., 2021; Wanner et al., 2021) indicate that NAS consistently enriches reference sequences (herein, mitochondrial sequence data) at levels ranging from 0.96-fold to 5.4-fold versus control ONT experiments. The NAS experiments conducted herein successfully captured entire mitogenomes of the hematophagous insects, with coverages ranging from 56x (*Ch. niger*) to 1,342x (*Cu. territans*). Moreover, the mitogenome of the control run (Exp. D; NAS off; *Cu. restuans*) was not fully assembled into a single, circular contig. It is possible that despite an estimated coverage of the mitochondrial genome (>29,000 sequences from the control experiment mapped to *Cu. pipiens* genome), there are likely nuclear DNA sequences containing pseudogenes that survived filtering, impacting the assembly of the mitochondrial genome and ultimately resulting in 19 short, linear contigs during the assembly process. Nuclear copies of mitochondrial DNA, or pseudogenes, have been reported in mosquitoes (Hlaing et al., 2009); and, NAS may resolve the issue of capturing nuclear pseudogenes often reported in traditional molecular methods by enriching for mitochondrial DNAs. Similar to Wanner et al. (2021), the results reported herein clearly document the utility of NAS for the enrichment and downstream *de novo* assembly of complete mitochondrial genomes. In light of these observations, we posit that NAS will be a particularly useful tool for a wide variety of studies, ranging from the inter- and intra-species characterization of mitogenome variation to biodiversity assessments of a wide variety of taxa wherein mitochondrial-based species barcoding is routinely conducted (Costa & Carvalho, 2010).

### 4.4 Species identification, biosurveillance, and educational potential

The taxonomic identification of hematophagous insects can be particularly challenging for both novice and skilled entomologists using morphological characters alone, especially for cryptic species. Moreover, given range expansions associated with climate change and anthropogenic-driven introductions, traditional morphological keys for a given geographic area may be unable to resolve species status for resident hematophagous insects. For these reasons, a robust molecular-based approach that facilitates the taxonomic identification of blood-feeding insects will greatly aid One Health initiatives rooted in biosurveillance.

The output of our NAS mitogenome sequencing approach consisted of enriched mitochondrial sequences ranging from hundreds to thousands of nucleotide bases of a given sample and downstream bioinformatic processes yielded *de novo* mitogenome assemblies for four insects for which no previous mitogenome existed (i.e., unavailable on public nucleotide repositories). The vast majority of available sequences for mitochondrial-based molecular barcoding consist of COI sequences (~9.87M) managed by the Barcode of Life Data System (BOLD; www.boldsystems.org). Thus, when combined with the BOLD database, the NAS mitogenome sequencing approach is particularly useful for the molecular identification of insect vectors. Select mitochondrial genes that are frequently used for species barcoding (i.e., COI, cyt-b) are easily extracted from *de novo* assembled mitogenomes of hematophagous insects (and their corresponding bloodmeals), for rapid BLAST and phylogenetic analyses facilitated by the BOLD initiative. We anticipate that the availability of complete mitogenome assemblies will increase exponentially with the usage of single-molecule long-read sequencing methodologies (e.g., ONT and Pacific Biosciences). From a One Health perspective, reference mitogenomes of hematophagous insects will provide important insights into the evolutionary histories of disease vectors and will greatly assist with accurate taxonomic identification efforts of cryptic species. We also note the educational potential of NAS, as the method can be easily incorporated into science curriculum across many educational levels. We demonstrate this utility by summarizing the natural history of species identified herein (collected in Minnesota, USA), using a combination of NAS discovery and BOLD analyses:

- *Aedes vexans* is a cosmopolitan species and often represents the most common mosquito species collected in the Upper Midwest, USA (Dunphy, Rowley, & Bartholomay, 2014). It is considered a competent vector of a variety of pathogens including West Nile virus, Eastern and Western encephalitis virus, Zika virus, St. Louis encephalitis virus, Rift Valley fever virus, and *Dirofilaria immitis* (Gendernalik et al., 2017; Ndiaye et al., 2016; Turell et al., 2005). Despite this, it is rarely considered an arthropod of significant disease concern. The mitochondrial genome assembly provided herein and deposited on NCBI (accession number OL351547) is the first reported for this species.
- *Aedes trivittatus* shares many ecological characteristics with *Ae. vexans* and is an abundant summer floodwater mosquito throughout the midwestern United States. *Ae. trivittatus* appears to feed chiefly on mammals, including the eastern cottontail rabbit (*Sylvilagus floridanus*), (Molaei, Andreadis, Armstrong, & Diuk-Wasser, 2008; Nasci, 1984; Pinger & Rowley, 1975), a feeding preference which is supported by our molecular findings. Like *Ae. vexans*, it is likely a competent vector of both West Nile virus (Tiawsirisup, Platt, Evans, & Rowley, 2005) and *Dirofilaria immitis* (Pinger, 1982), though its precise role in pathogen transmission in North America has not been well-characterized. The mitochondrial genome assembly provided herein and deposited on NCBI (accession number PENDING) is the first reported for this species.
- *Culex restuans* is distributed across most of North America from Guatemala north throughout the US and Canada (Darsie & Ward, 2005; Strickman & Darsie, 1988). It is being increasingly recognized as an important vector of West Nile virus, particularly in urban and suburban areas (Johnson, Robson, & Fonseca, 2015), and is likely capable of transmitting St. Louis encephalitis virus (Monath & Tsai, 1987). Notably, adult female *Cu. restuans* mosquitoes are largely indistinguishable from North America's primary West Nile virus vector, *Cu. pipiens*, based on external morphologic features (Apperson et al., 2002; Ebel, Rochlin, Longacker, & Kramer, 2005). The mitochondrial genome assembly provided herein and deposited on NCBI (accession number OL351548) is the first reported for this species and represents a key molecular resource for the molecular differentiation of these medically-important West Nile virus vectors.
- *Culex territans* is widespread in the Eastern US where it feeds on frogs (Bartlett-Healy, Crans, & Gaugler, 2008). It transmits a hepatozoon parasite to frogs but is unlikely to be involved in mammalian disease transmission (Desser, Hong, & Martin, 1995). The mitochondrial genome assembly provided herein and deposited on NCBI (accession number OL351549) is the first reported for this species.
- *Chrysops niger* is widely distributed in the Eastern US and SE Canada and is associated with marshy areas where its larvae feed on organic matter in soil (Pechumann, 1973). It is a common pest of livestock (Weiner & Hansens, 1975). Other members of the genus are involved in the mechanical transmission of *Fransicella tularensis* in the Western US (Jellison, 1950). The mitochondrial genome assembly provided herein and deposited on NCBI (accession number OL351550) is the first reported for this species.

## 5. Conclusions

The ONT NAS method is a revolutionary approach that can be leveraged to address many biological questions and is particularly useful for zoonotic pathogen surveillance (Kipp et al., 2021). When combined with bioinformatic depletion of host DNA/RNA, it is likely that NAS will enhance pathogen discovery within a wide-variety of vector species. Long-read single-molecule sequencing is ideally suited for recovering entire mitogenomes across the tree of life, thus opening the door to enhanced mitochondrial barcoding. Future work by our team will focus on leveraging NAS for metagenomic ‘xenosurveillance’ approaches aimed at identifying both human and animal viral pathogens associated with mosquito bloodmeals in high-risk tropical regions (Grubaugh et al., 2015) as well as to identify hematophagous insects associated with poorly understood vector-borne diseases of livestock and wildlife. Given the well-documented ease of use and portability of the ONT MinION (Blanco et al., 2020; Maestri et al., 2019; Pomerantz et al., 2018), molecular experiments can be performed within individual research laboratories, including field-based locations, with results obtained in minutes or hours. Thus, MinION NAS-based surveillance of hematophagous insects would effectively inform One Health research projects and biosurveillance initiatives in real-time.

## Data accessibility

All raw sequence data generated during the course of this study have been deposited within the NCBI SRA biorepository and are publicly available (Bioproject: PRJNA775614; Biosamples: SAMN22604479-SAMN22604483; SAMN22888850-SAMN22888851). Mitochondrial genome assemblies resulting from our research have been deposited at GenBank (NCBI Accession OL351547-OL351550).

## Authors’ contributions

PAL and JO envisioned the research. EK, LLL, CB, JB performed fieldwork. EK, LLL, CB, JB, MSM generated molecular data. EK, LLL, CF, PAL conducted bioinformatic analyses and interpreted the results. All authors contributed to the writing of the initial draft and all edited and approved the final manuscript.

## Acknowledgements

We thank Suzanne Stone for assistance within the molecular laboratory and related logistics. The Minnesota Supercomputing Institute provided essential computational and data storage resources. Staff of the Wildlife Rehabilitation Center of Minnesota (Roseville, MN) and Luciano Caixeta kindly provided access to forested lands and the University of Minnesota dairy barn, respectively, for insect collecting. Financial support for CMB and JB was kindly provided by the University of Minnesota, College of Veterinary Medicine Summer Scholars program with funds originating from the Office Of The Director of the NIH under Award Number T35OD011118 and the Office of Graduate Programs, College of Veterinary Medicine. We thank Tiffany Wolf and Erin Burton for providing dissecting microscopes for morphological identifications of insects. We used BioRender (www.biorender.com) to create figures presented herein.

