## Supplemental Figure 1 for "Nanopore adaptive sampling for mitogenome sequencing and bloodmeal identification in hematophagous insects"

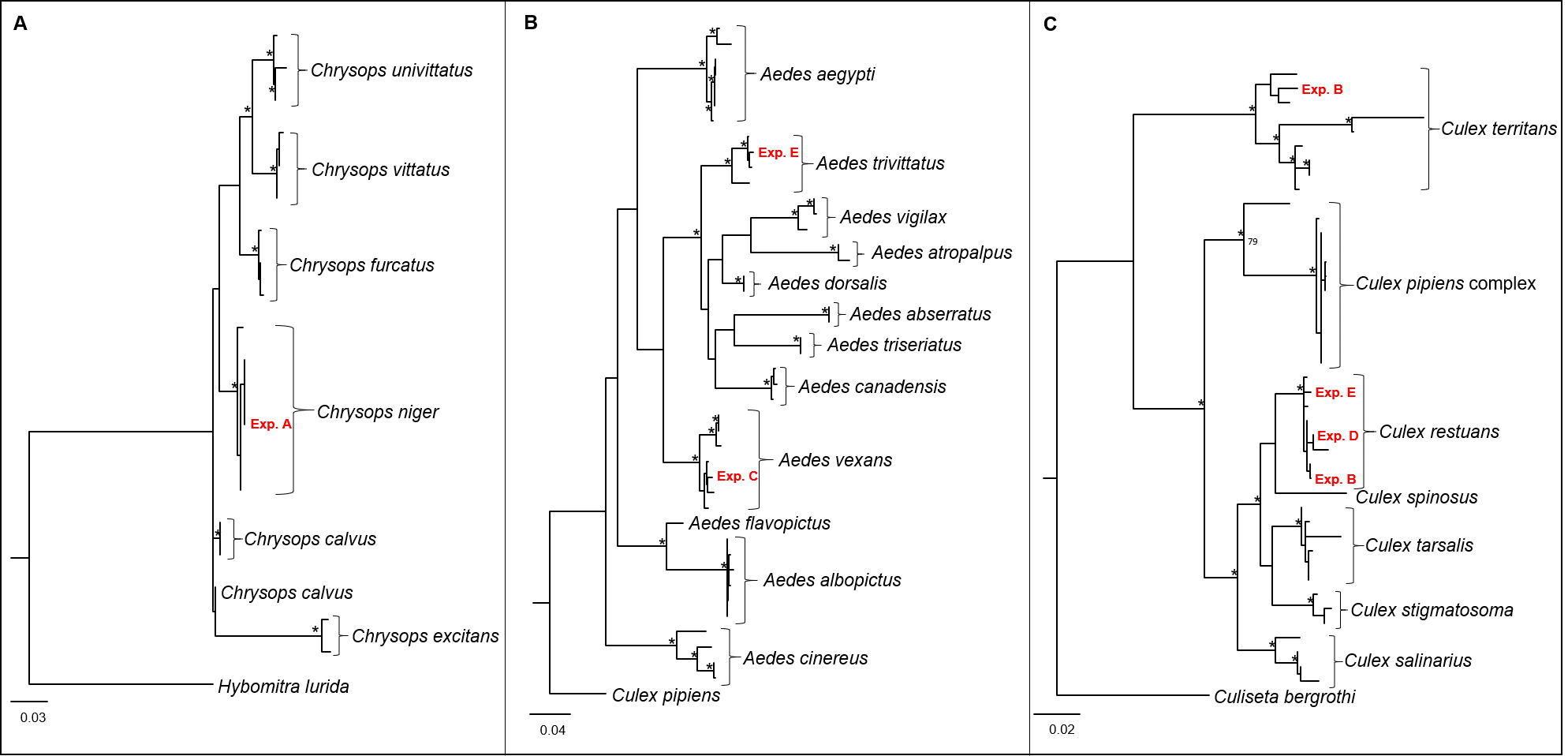

**Supplemental Figure 1. Phylogenetic trees constructed of vector species (*Chrysops*, *Aedes*, and *Culex*) COI sequences.** Barcoded sequences generated from each experiment (Exp. A-E) are denoted in red and statistically supported nodes (≥ 75) are denoted with a single asterisk (*). **A:** Maximum likelihood analysis with 1,000 bootstrap replicates for COI of *Chrysops* flies with *Hybomitra* *lurida* used as an outgroup. **B:** Maximum likelihood analysis with 1,000 bootstrap replicates for COI of *Aedes* mosquitoes with *Culex* *pipiens* used as an outgroup. **C:** Maximum likelihood analysis with 1,000 bootstrap replicates for COI of *Culex* mosquitoes with *Culiseta* *bergrothi* as an outgroup.
