## Supplemental Table 1 for "Nanopore adaptive sampling for mitogenome sequencing and bloodmeal identification in hematophagous insects"

**Supplemental Table 1.** Vector specimens examined for each sequencing experiment. SRA Accession numbers are reported for raw fastq files from each experiment. NCBI Organelle Accession Numbers are only available for the four vector species that had fully assembled mitochondrial genomes generated in this study.

| **Experiment ID** | **Species** | **Collection Date** | **Locality** | **GPS Coordinates** | **SRA Accession Number** | **NCBI Organelle Accession Number** |
| --- | --- | --- | --- | --- | --- | --- |
| A | *Chrysops niger* | 07 June 2021 | USA: Minnesota; Washington Co. | 44.860946 N 92.780562 W | SAMN22604483 | OL351550 |
| B | *Culex restuans* | 02 June 2021 | USA: Minnesota; Ramsey Co. | 44.975747 N 93.190584 W | SAMN22604480 | OL351458 |
| B | *Culex territans* | 02 June 2021 | USA: Minnesota; Ramsey Co. | 44.975747 N 93.190584 W | SAMN22604482 | OL351549 |
| C | *Aedes vexans* | 02 June 2021 | USA: Minnesota; Ramsey Co. | 44.975747 N 93.190584 W | SAMN22604479 | OL351547 |
| D | *Culex restuans* | 02 June 2021 | USA: Minnesota; Ramsey Co. | 44.975747 N 93.190584 W | SAMN22604481 | N/A |
| E | *Culex restuans* | 02 June 2021 | USA: Minnesota; Ramsey Co. | 44.975747 N 93.190584 W | SAMN22888850 | N/A |
| E | *Aedes trivittatus* | 02 June 2021 | USA: Minnesota; Ramsey Co. | 44.975839 N 93.190618 W | SAMN22888851 | N/A |
