## Supplemental Table 2 for "Nanopore adaptive sampling for mitogenome sequencing and bloodmeal identification in hematophagous insects"

**Supplemental Table 2.** Genbank accession numbers that were included in the reference fasta file used during NAS sequencing runs. Exp. A included a closely related fly species and possible bloodmeals (a vector species not used in this study was included in this reference file). Exp. B contained a closely related mosquito vector species, possible bird and mammal bloodmeals, and possible pathogen targets of interest. Exp. E contained closely related vector species and possible bird, mammal, reptile, and amphibian bloodmeals. Exp. C was not included in this table because we used the RefSeq download for the entire mitochondrial genome database provided by NCBI (<https://ftp.ncbi.nlm.nih.gov/refseq/release/mitochondrion/>; accessed on 16 September 2021).

| **Exp. A** | **Exp. B** | **Exp. E** |
| --- | --- | --- |
| *Bos taurus*, mitochondrion: AF492351.1 | *Canis lupus familiaris*, mitochondrion: NC_002008.4 | *Canis lupus familiaris,* mitochondrion: NC_002008.4 |
| *Ovis aries*, mitochondrion: NC_001941.1 | *Sciurus carolinensis*, mitochondrion: NC_050012.1 | *Sciurus carolinensis*, mitochondrion: NC_050012.1 |
| *Equus caballus*, mitochondrion: NC_001640.1 | *Procyon lotor*, mitochondrion: AB297804.1 | *Procyon lotor*, mitochondrion: AB297804.1 |
| *Columba livia*, mitochondrion: NC_013978.1 | *Sylvilagus floridanus*, mitochondrial genes: MW323423.1, MK967948.1, KU057246.1, AY011158.1, KC923400.1 | *Sylvilagus floridanus*, mitochondrial genes: MW323423.1, MK967948.1, KU057246.1, AY011158.1, KC923400.1 |
| *Culicoides arakawae*, mitochondrion: AB361004.1 | *Peromyscus leucopus*, mitochondrion: NC_037180.1 | *Peromyscus leucopus*, mitochondrion: NC_037180.1 |
| *Homo sapiens*, mitochondrion: NC_012920.1 | *Homo sapiens*, mitochondrion: NC_012920.1 | *Homo sapiens*, mitochondrion: NC_012920.1 |
| *Culicoides crepuscularis* COI gene: KR689448.1 | *Didelphis virginiana*, mitochondrion: Z29573.1 | *Didelphis virginiana*, mitochondrion: Z29573.1 |
| *Chrysops silvifacies*, mitochondrion: KT225292.1 | *Odocoileus virginianus*, mitochondrion: NC_015247.1 | *Odocoileus virginianus*, mitochondrion: NC_015247.1 |
| *Francisella tularensis*, genome: CP073120.1 | *Eptesicus fuscus*, mitochondrion: MF143474.1 | *Eptesicus fuscus*, mitochondrion: MF143474.1 |
| -- | *Felis catus*, mitochondrion: NC_001700.1 | *Felis catus*, mitochondrion: NC_001700.1 |
| -- | *Rattus norvegicus*, mitochondrion: AY172581.1 | *Cardinalis cardinalis*, mitochondrion KM078795.1 |
| -- | *Mus musculus*, mitochondrion: KY018919.1 | *Haemorhous mexicanus*, mitochondrion: KM078782.1 |
| -- | *Microtus ochrogaster*, mitochondrion: KT166982.1 | *Passer domesticus*, mitochondrion: KM078784.1 |
| -- | *Vulpes vulpes*, mitochondrion: AM181037.1 | *Sturnus vulgaris*, mitochondrion: KT946692.1 |
| -- | *Canis latrans*, mitochondrion: DQ480509.1 | *Corvus brachyrhynchos ,* mitochondrion: KP403809.1 |
| -- | *Blarina brevicauda*, mitochondrion: NC_042734.1 | *Turdus migratorius*, mitochondrion: KJ909198.1 |
| -- | *Buteo jamaicensis*, mitochondrial genes: AY987334.1, DQ434504.1, U83720.2, HM454161.1 | *Branta canadensis*, mitochondrion: DQ019124.1 |
| -- | *Strix varia*, mitochondrion: MF431745.1 | *Columba livia*, mitochondrion: NC_013978.1 |
| -- | *Cardinalis cardinalis*, mitochondrion KM078795.1 | *Anas platyrhynchos*, mitochondrion: NC_009684.1 |
| -- | *Haemorhous mexicanus*, mitochondrion: KM078782.1 | *Meleagris gallopavo*, mitochondrion: JF275060.1 |
| -- | *Passer domesticus*, mitochondrion: KM078784.1 | *Chrysemys picta*, mitochondrion: NC_002073.3 |
| -- | *Sturnus vulgaris*, mitochondrion: KT946692.1 | *Chelydra serpentina*, mitochondrion: EF122793.1 |
| -- | *Corvus brachyrhynchos ,* mitochondrion: KP403809.1 | *Storeria dekayi*, mitochondrial genes: MH274692.1, AF471050.1, AF402639.1 |
| -- | *Turdus migratorius*, mitochondrion: KJ909198.1 | *Ambystoma tigrinum*, mitochondrion: AY659992.1 |
| -- | *Branta canadensis*, mitochondrion: DQ019124.1 | *Anaxyrus americanus*, mitochondrion: NC_047224.1 |
| -- | *Columba livia*, mitochondrion: NC_013978.1 | *Aedes albopictus*, mitochondrion: NC_006817.1 |
| -- | *Anas platyrhynchos*, mitochondrion: NC_009684.1 | *Culex pipiens*, mitochondrion: HQ724614.1 |
| -- | *Meleagris gallopavo*, mitochondrion: JF275060.1 | -- |
| -- | *Plasmodium gallinaceum*, mitochondrion: AB250690.1 | -- |
| -- | *Plasmodium relictum*, mitochondrion: AY733088.1 | -- |
| -- | *Dirofilaria immitis*, mitochondrion: AJ537512.1 | -- |
| -- | *Plasmodium relictum*, apicoplast: NC_031964.1 | -- |
| -- | *Plasmodium gallinaceum*, apicoplast: NC_031963.1 | -- |
| -- | *Leucocytozoon fringillinarum*, mitochondrion: FJ168564.1 | -- |
| -- | *Leucocytozoon majoris*, mitochondrion: FJ168563.1 | -- |
| -- | *Culex pipiens*, mitochondrion: HQ724614.1 | -- |
| -- | *Aedes albopictus*, mitochondrion: NC_006817.1 | -- |
