## Supplemental Table 3 for "Nanopore adaptive sampling for mitogenome sequencing and bloodmeal identification in hematophagous insects"

**Supplemental Table 3.** Sequences were randomly subsampled starting at 9,000 sequences. The resulting subsampled files were mapped to the mitogenome of *Culex* *pipiens* using minimap2.

|  | **Randomly subsampled number of sequences** | | | | |  |
| --- | --- | --- | --- | --- | --- | --- |
| **Sample** | **9,000** | **5,000** | **2,000** | **1,000** | **500** | **Avg Mapped (%)** |
| D: *Culex restuans* | 172 | 96 | 34 | 23 | 12 | 2.04 |
| B: *Culex restuans* | 6,747 | 3,723 | 1,496 | 737 | 366 | 74.2 |
| B: *Culex territans* | 5,875 | 3,252 | 1,301 | 655 | 336 | 65.6 |
