## Supplemental Table 4 for "Nanopore adaptive sampling for mitogenome sequencing and bloodmeal identification in hematophagous insects"

**Supplemental Table 4.** Sequences downloaded from GenBank (species and accession numbers) used in the phylogenies of bloodmeals sequenced from Exp. E.

| **House Sparrow** | **Cottontail** |
| --- | --- |
| *Passer domesticus:* MN356394.1, KM078784.1, AF407128.1 | *Sylvilagus floridanus:* AY012126.1, KU057246.1 |
| *Passer montanus:* MH211396.1, KM577704.1 | *Sylvilagus audobonii:* U67285.1 |
| *Passer ammodendri*: KT895996.1 | *Sylvilagus nuttallii*: KU057255.1 |
| *Montifringilla taczanowskii:* KJ148631.1 | *Lepus californicus:* KJ397614.1 |
| -- | *Lepus americanus:* KJ397613.1 |
